# Phosphatidylinositol 4-kinase II β downregulation induces ER stress mediated ROS generation and increases radio-sensitivity in MCF-7

**DOI:** 10.1101/2023.10.03.557937

**Authors:** Sonica Chaudhry, Rahul Checker, Maikho Thoh, Santosh Kumar Sandur, Gosukonda Subrahmanyam

## Abstract

Type II phosphatidylinositol 4-kinase has historically been associated with vesicular trafficking and growth factor receptor signaling. Recently, it was shown to influence apoptosis, cell adhesion, motility, and inflammatory reactions. Previous study from our group showed that down regulation of PtdIns 4-kinase II β inhibits NF-κB translocation to the nucleus and induce apoptosis in cancer cell lines. Since, NF-κB regulates redox homeostasis and stress signaling, the role of PtdIns 4-kinase II β in regulating these signaling cascades was investigated using human breast cancer cell line (MCF-7 cells). Knockdown of PtdIns 4-kinase II β inhibited NF-κB nuclear translocation and down regulated Bcl-2 and AKT-3 mRNA levels. Silencing of PtdIns 4-kinase II β increased ROS levels with a concomitant decrease in GSH: GSSG ratio, increased DNA damage and induced apoptosis. Inhibition of NADPH oxidase (NOX) activity restored redox homeostasis. Since NOX gets activated during ER stress and the resultant increase in ROS can induce apoptosis, we studied the role of PtdIns 4-kinase II β shRNA in ER stress and observed an increase in ER stress markers, pERK, IRE-1α, BiP and PDI and up regulation of GADD153. At cellular level, these cells showed ER deformity. ROS scavengers and ER chemical chaperone could rescue MCF-7 cells from PtdIns 4-kinase II β shRNA induced apoptosis. These results indicate involvement of PtdIns4-kinase II β in regulation of ER function. Interestingly, irradiation of MCF-7 cells increased PtdIns 4-kinase II β mRNA levels and knock-down of PtdIns 4-kinase II β increased radio-sensitivity of the cells indicating potential role of PtdIns 4-kinase II β in cancer radio-resistance.

## Introduction

PtdIns 4-kinase(s) are known to regulate cell proliferation through PtdIns 3-kinase and Phospholipase C signaling pathways. Among all the isoforms of PtdIns 4-kinase(s), the physiological significance of PtdIns 4-kinase II β is least understood. PtdIns 4-kinase II β is localized to membranes and cytosol ^1^. At plasma membrane, PtdIns 4-kinase II β has been shown to associate with growth factor receptors such as EGF receptor, TCR-CD3 complex and FcεR1 ^2–4^. Cytosolic fraction of PtdIns 4-kinase II β has low enzymatic activity and its role in signaling is not well understood. Structural insights of PtdIns 4-kinase II β shows presence of intrinsically disorganized regions, PXXP domains, SH3 binding domains, and atypical leucine zipper region that may help in protein-protein interactions. In human, PtdIns 4-kinase II β gene is located on chromosome 4p15.2. In addition to PtdIns4-kinase II β, this region contains putative tumor suppressor genes ^5^. Heterozygous loss of PtdIns 4-kinase II β is shown to be associated with lung (squamous cell and adenocarcinoma), esophageal, pancreatic, prostate, breast, liver, and various other epithelial cancers ^6^. PtdIns 4-kinase II β was shown to associate with a tumor suppressor, Prostate apoptosis response-4 (Par-4). The depletion of PtdIns 4-kinase II β was shown to increase half life of Par-4 protein and facilitates its nuclear localization ^7^.Recently, kinome expression profiling identified PtdIns 4-kinase II β gene associated with poor prognosis in multiple myeloma patients ^8^.

Reactive oxygen species (ROS) are oxygen derived molecules that have unpaired electron(s) in their outermost electronic shell ^9^. Intracellularly, ROS are generated as metabolic by-products of oxidative phosphorylation in mitochondria. Additionally, peroxisomal activity, Cyt P450, cyclooxygenases, lipoxygenases, and thymidine phosphorylase also generate ROS as byproducts ^9^. In contrast, NADPH oxidases produce ROS as part of their biological functions ^10^. ROS oxidize various biological molecules including nucleic acids, proteins and lipids. ROS react with nucleic acids and induce single and double stranded DNA breaks. The imbalance between ROS production and ROSscavenging leads to oxidative stress ^11^.In mitochondria, membrane permeability transition pore (MPTP) proteins get oxidized by ROS which results in MPTP opening and release of cytochrome c. The multimeric complex of Apoptotic protease activating factor-1 (Apaf-1), cytochrome c and caspase 9 activates caspase cascade leading to apoptosis ^12^.Cells have developed various defense mechanisms to combat oxidative stress. NF-κB, a transcriptional factor, upregulates the expression of anti-oxidant proteins such as manganese superoxide dismutase, ferritin heavy chain, *thioredoxins*, glutathione S-transferase, NAD(P)H dehydrogenase [quinone] 1 and glutathione peroxidase-1, and dihydrodiol dehydrogenase ^13^. These proteins are involved in alleviating oxidative stress and promote cell survival.Cancer cells are more susceptible to apoptotic induction by radiation and other chemotherapeutic drugs when subjected to oxidative stress as compared to normal cells ^14^. The combinatorial radiotherapy with drugs promoting oxidative stress may help in better cancer treatment.

PtdIns(4)*P* is suggested to play an important role in ER and plasma membrane (PM) crosstalk^15^. ER and PM forms a close connection with each other without undergoing membrane fusion, these connecting points is called ER-PM junctions. These junctions play a major role in maintaining ER structure and functions. Loss of tethering proteins at these junctions can lead to ER stress ^16^. PtdIns(4,5)*P*_*2*_ is suggested to regulate levels of phosphatidylserine and PtdIns in plasma membrane ER-PM junctions ^17^. The isoform specific role of PtdIns 4-kinase(s) in ER stress pathways are not clearly understood. Recently, PtdIns*4P* was shown to regulate mitochondrial fission. The depletion of type III PtdIns 4-kinase induced mitochondrial hyper fusion and branching. The group also reported that depletion of PtdIns 4kinase II β resulted in mild hyper fusion of mitochondria in Hela cells^18^.

In the present study we have explored the role of PtdIns 4-kinase II β on stress pathways with the aim that ER stress associated ROS generation may sensitize cancer cells to radiation induced killing.

## MATERIAL AND METHODS

### Materials

MCF-7 cell line was procured from National Centre for Cell Science, Pune, India. PtdIns 4-kinase II β shRNA and scrambled shRNA sequences cloned in pmu6 vector as described earlier. Lipofectamine 3000 was from Invitrogen, Thermo Fisher Scientific, USA. Diphenyleneiodonium, TroloX, Phenyl butyric acid, PEG-catalase, PEG-superoxide dismutase, N-acetyl cysteine (NAC), 5,5’-dithio-bis(2-nitrobenzoic acid) (DTNB), Glutathione reductase (GR), NADPH, vinyl pyridine and triethanolamine were from Sigma-Aldrich. cmDCFDA (chloromethyl derivative of dihydrodichloroflourescin diacetate) was from Life Technologies, Thermo Fisher Scientific, USA. Anti type II PtdIns 4-kinase β (ab37812) were from Abcam, UK. Caspase 8 and caspase 9 inhibitors were from Calbiochem, Merck, Germany. Caspase 12 inhibitor was from Abcam, UK. Anti β-actin (#4970), anti HA (#3724), anti Myc (#2278), anti BiP (#3177), anti calnexin (#2679), anti PDI (Protein disulphide isomerase) (#3501), anti Ero1-Lα (#3264), anti IRE1α (#3294) and anti CHOP (#2895) antibodies were from Cell Signaling Technology, USA. Electron microscopy grade osmium tetraoxide and glutaraldehyde were from Sigma-Aldrich. Epoxy resins for embedding transmission electron microscope samples were from Toshniwal Brothers Pvt Ltd, Bangalore, India.

### Cell culturing and transfection

MCF-7 cells were from National Centre for Cell Science, Pune, India. MCF-7 cells were grown in Dulbecco’s Modified Eagle’s Medium (DMEM) with 10% fetal bovine serum at 37ºC with 5% CO_2_ in a humidified incubator.Cells were transfected using Lipofectamine 3000 transfection reagent in antibiotic free media as per manufacturer’s protocol.

### Apoptosis assay by Propidium Iodide staining (cell cycle analysis)

MCF-7 cells were transfected with either control or PtdIns 4-kinase II β shRNA plasmids and incubated for 36 h. Cells were harvested and fixed using ice cold 70% ethanol at - 20ºC overnight. Cells were washed with PBS three times followed by propidium iodide staining (50 µg/mL propidium iodide, 50 µg/mL RNase in PBS, 0.1% sodium citrate, 0.1% Triton X-100) on ice for 30 min. Flow cytometric quantification of propidium iodide stained cells were carried out for 20,000 cells using Partec flow cytometer and analysed using Flowjo software ^19^

MCF-7 cells were transfected with either control or PtdIns 4-kinase II β shRNA plasmids and incubated for 24 h. Cells were harvested and washed with 0.5 % BSA in PBS three times. Cells were resuspended in binding buffer (5mM HEPES, 70mM NaCl and 1.25mM CaCl_2_ at pH 7.4) followed by staining with annexinV as per manufacturer’s protocol. Cells were incubated in dark for 20 min. Propidium iodide was added to cells followed by immediate analysis on flow cytometer. Quantification was done for 20,000 cells using Partec flow cytometer and analysed using Flowjo software.

### ROS estimation

MCF-7 cells were pre incubated with CM-H_2_DCFDA (10 µM) for 1 h. These cells were transfected with either control or PtdIns 4-kinase II β shRNA plasmids and cultured for 24 h. Fluorescence was measured using a Microplate reader (excitation wavelength 485 nm, emission wavelength 535 nm and Gain 50).

MCF-7 cells were pre incubated with CM-H_2_DCFDA (10 µM) for 1 h. These cells were transfected with either control or PtdIns 4-kinase II β shRNA plasmids. After 4 h of transfection, diphenyleneiodonium chloride (DPI) (50 µM) was added to the cells. Cells were cultured for 36 h and fluorescence was measured using a Microplate reader.

### Treatment with ROS scavenger

MCF-7 cells were pre-incubated with NAC (10 mM), Trolox (1 mM), PEG-superoxide dismutase (135 U/mL) and PEG-catalase (200 U/mL) for 1 hprior to transfection. Cells were transfected with either control or PtdIns 4-kinase II β shRNA plasmids and incubated for 36 h. Apoptotic cells were analyzed on flow cytometer using propidium iodide as described above.

### GSH: GSSG estimation

MCF-7 cells were transfected with either control or PtdIns 4-kinase II β shRNA plasmids and incubated for 24 h. Cells were harvested and lysed in extraction buffer (0.1% Triton X-100, 0.6% sulfosalicylic acid, 5 mM EDTA, 10 mM potassium phosphate buffer). Cells were lysed by sonication and freeze thaw cycle (repeated three times). Lysates were centrifuged at 12000 ***g*** for 10 min. Supernatants were collected and distributed in two tubes. For GSH estimation, 20 µL cell lysate was added to 120 µL of DTNB (150 mM) and GR (10 U/mL) mix (1:1) and incubated at 37°C for 60 min.Reaction was terminated by adding 60 µL of NADPH (0.8 mM) solution and absorption was measured at 412 nm. For GSSG estimation, GSH was scavenged by adding 2 µL of vinyl pyridine to 100 µL of lysate and incubated at 37°C for 60 min. Subsequently, 6 µL of triethanolamine was added to the side of tube with vigorous mixing and incubated for 10 min. Lysates were then processed for GSH estimation as described above. Total GSH and GSSG ratios were calculated as described earlier ^20^.

### Single cell electrophoresis

MCF-7 cells were transfected with either control or PtdIns 4-kinase II β shRNA plasmids and incubated for 24 h. Cells were harvested and mixed with 1.5 mL of 0.8% low melting agarose solution uniformly layered on frosted slides. After solidification, slides were kept in 50 mL lysis buffer (2.5 M NaCl, 100 mM Na-EDTA with freshly added 1% Triton X-100 and 10% DMSO) at 4 □C for 1 h. Slides were washed with 0.5 X TBE (45 mM Tris-Borate (pH 8.3), 1 mM EDTA) and incubated in same buffer at room temperature for 20 min. Electrophoresis was carried out for 20 min at 1.1 V/cm. Slides were stained with SYBR green and visualized at 40X magnification using fluorescent microscope (Axioplan, Carl-Zeiss Germany). In each group, 100 cells were imaged and analyzed using CASP software ^19,21^.

### Electrophoretic Mobility Shift Assay

MCF-7 cells were transfected with either control or PtdIns 4-kinase II β shRNA plasmids and incubated for 24 h. These transfected cells were suspended in cytoplasmic extraction buffer (10 mM HEPES (pH 7.4), 10 mM KCl, 10 mM EDTA, 5 mM EGTA) with intermittent vortexing on ice for 1 h. The cytoplasmic extract was centrifuged at 15000 ***g*** for 3 min. Nuclear pellet was suspended in nuclear extraction buffer (5 mM HEPES (pH 7.4), 1.5 mM KCl, 4.6 M NaCl, 10 mM EDTA, 5 mM EGTA and 20% glycerol) with intermittent vortexing for 3 h.The nuclear extract was collected by centrifugation at 15000 ***g*** for 30 min. Nuclear extract (8 μg) was added to ^32^P (16 *f*mol) end labelled NF-κBoligo (5’-TTGTTACAAGGGACTTTCCGCTGGGGACTTTCCAGGGAGG CGTGG3’),

0.5 □g poly (2□-deoxyinosinic-2□-deoxycytidylic acid) in binding buffer (25 mM HEPES (pH 7.9), 150 mM NaCl, 0.5 mM EDTA, 0.5 mM dithiothreitol, 1% Nonidet P-40, 5% glycerol). The above mixture was incubated at 37ºC for 1 h. Native polyacrylamide gel (6.6%) electrophoresis was carried out in running buffer (50 mM Tris (pH 8.5), 200 mM Glycine, 1 mM EDTA). The gel was dried and exposed on a Molecular-Dynamics PhosphorImager Screen. The Screen was scanned using a phosphorImager scanner (Amersham) ^22^.

### Transmission Electron Microscopy

MCF-7 cells were transfected with either control or PtdIns 4-kinase II β shRNA plasmids and incubated for 36 h. Cells were trypsinized and centrifuged at 1000 ***g***for 5 min. Cells were washed with sodium cacodylate (150 mM) and fixed by 0.2 % glutaraldehyde and 4% formaldehyde in sodium cacodylate (150 mM) at 4° C overnight. Fixed cells were washed twice with sodium cacodylate (150 mM). Cells were again fixed with osmium tetraoxide for 40 min and washed twice with sodium cacodylate (150 mM). Cell pellets were stained with uranyl acetate for 30 min. Cells were washed twice with distilled water. Cells were gradually dehydrated by washing with 30%, 50%, 70%, 95% and 100% ethanol for 10 min. Cells were incubated with increasing concentration of propidium oxide (25%, 50%, 90% and 100%) in ethanol. Cells were incubated at each concentration of propidium oxide on rotary mixer for 10 min. Resin was prepared fresh as per manufacturer’s protocol. Cells were incubated with increased gradient of epoxy resin (resin : propidium oxide :: 1:3, 1:1 and 3:1) on rotary mixer for 2 h. Cells were incubated with 100% resin overnight and cured at 80 °C for 48 h. Ultra-thin sections (70-100 nm) were obtained from microtome and stained with uranyl acetate and lead citrate. Transmission electron micrographs were obtained.

### Clonogenic assay

MCF-7 cells were transfected with either control or PtdIns 4-kinase II β shRNA plasmids. Sub optimal concentration of plasmids was used to reduce cell death in unirradiated PtdIns 4-kinase II β shRNA transfectants. After 12 h of incubation, 4000 cells were plated and irradiated with 2 Gy (gray), 4 Gy or 6 Gy doses of γ-radiation using a Co^60^gamma-source (Bhabhatron, BARC). Cells were incubated for 12 days for visible colony formation, stained with crystal violet and counted.

## RESULTS

### Depletion of PtdIns 4-kinase II β dysregulates redox homeostasis and induces DNA double strand breaks

Earlier studies from our lab have shown that knockdown of PtdIns 4-kinase II β correlated with increased apoptosis in MCF-7 cells ^7^. Induction of apoptosis was further confirmed by Annexin V staining in PtdIns 4-kinase II β knockdown MCF-7 cells (Fig. 1A). To determine the cause of apoptosis induction, we measured ROS levels in PtdIns 4-kinase II β shRNA plasmid transfected MCF-7 cells using DCF fluorescence. These cells showed an increase in ROS levels in comparison to control cells (Fig. 1B). Conversely, PtdIns 4-kinase II β depletion also resulted in reduction of GSH levels and an increase in its oxidised form, GSSG. The ratio of GSH and GSSG was reduced to 6:1 from 11:1 in PtdIns 4-kinase II β shRNA transfectants within 24 h (Fig. 1C). These results indicated that PtdIns 4-kinase II β depletion induced oxidative stress in these cells.

**Fig. 1.**
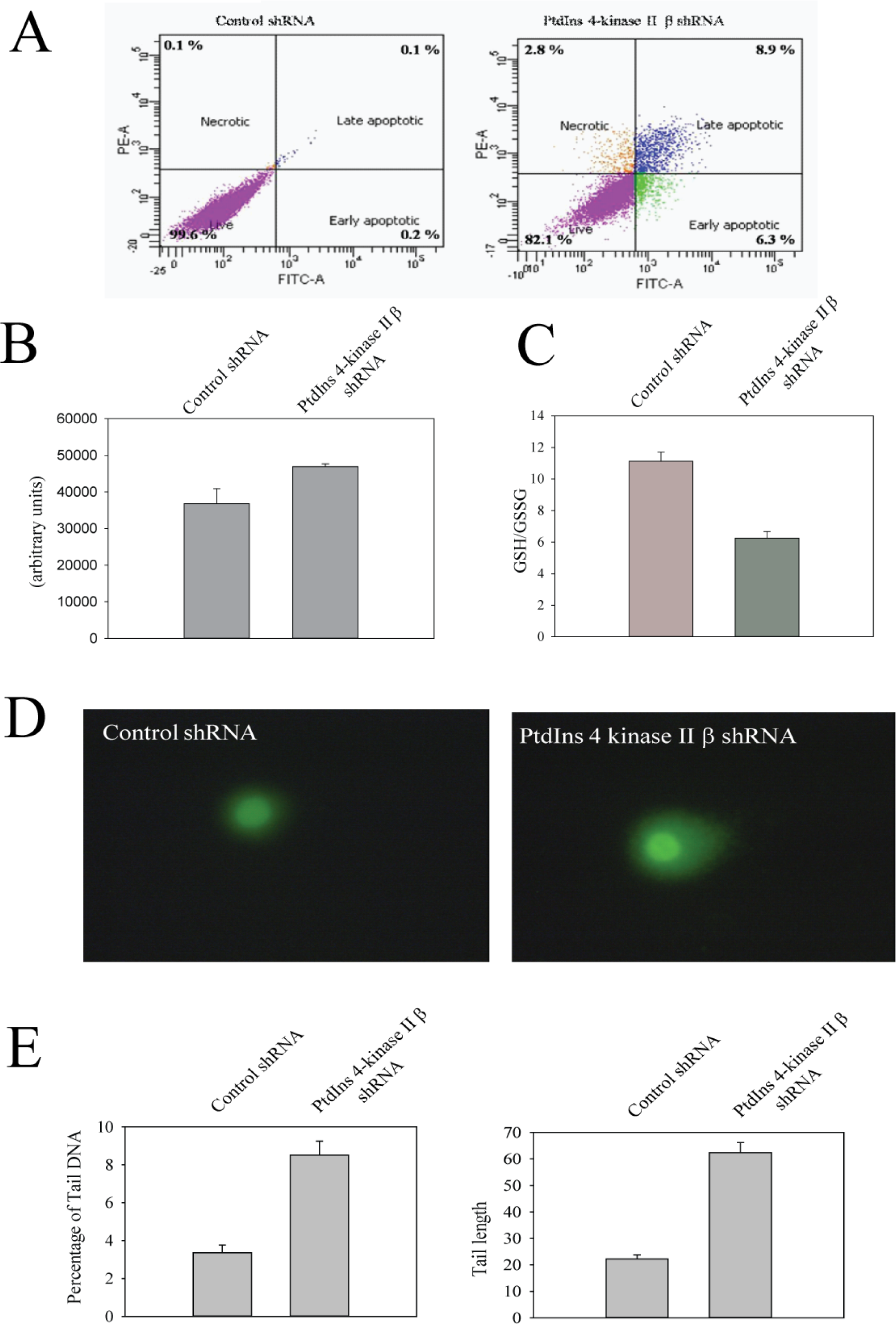
PtdIns 4-kinase II β shRNA induces oxidative stress in MCF-7 cells. MCF-7 cells were transfected with either control or PtdIns 4-kinase II β shRNA plasmids and incubated for 24 h. **A**.Cells were stained with FITC conjugated Annexin V and propidium iodide followed by flow cytometric analysis. Numbers in quadrant depict percentage of cells. **B**.Transfected cells were pre-incubated with CM-H_2_DCFDA for 1 h. After 24 h of transfection, fluorescence was measured (excitation: 485 nm, emission: 528 nm and gain: auto scale)using microplate reader. **C**.The cytoplasmic extracts were assayed for GSH and GSSG as described in Materials and Methods. The ratio of GSH and GSSG in each sample was calculated. **D**.The transfectants were subjected to single cell electrophoresis. Representative fluorescent images.**E**.Quantification of data. 100 cells were imaged for each group and images were analyzed with CASP software. Mean percentage of tail DNA and tail length were shown. (The data indicates mean ± SD. ** P<0.001). The experiments were repeated three times.

A consequence of increased ROS levels is oxidation of thiol groups, metal prosthetic groups and DNA damage. The effect of PtdIns 4-kinase II β depletion on DNA integrity was studied using single cell electrophoresis performed at neutral pH. The results showed an increase in percentage of tail DNA and tail length which implies double stranded DNA damage in PtdIns 4-kinase II β depleted cells (Fig. 1D & E). These results are consistent with the observation that PtdIns 4-kinase II β depletion leads to oxidative stress, DNA damage and apoptosis.

### ROS produced through activation of NOX

ROS are generated as by-products from mitochondria, peroxisomes and cyt P450. In addition, NADPH oxidases produce ROS as part of their biological functions. Inhibition of NADPH oxidase activity with diphenyleneiodinium abrogated increased ROS production in PtdIns 4-kinase II β depleted MCF-7 cells (Fig. 2A). JC1 staining showedincrease inmitochondrial membranepermeability indicating mitochondrial damage in PtdIns 4-kinase II β depleted cells (Fig. 2B).

**Figure 2.**
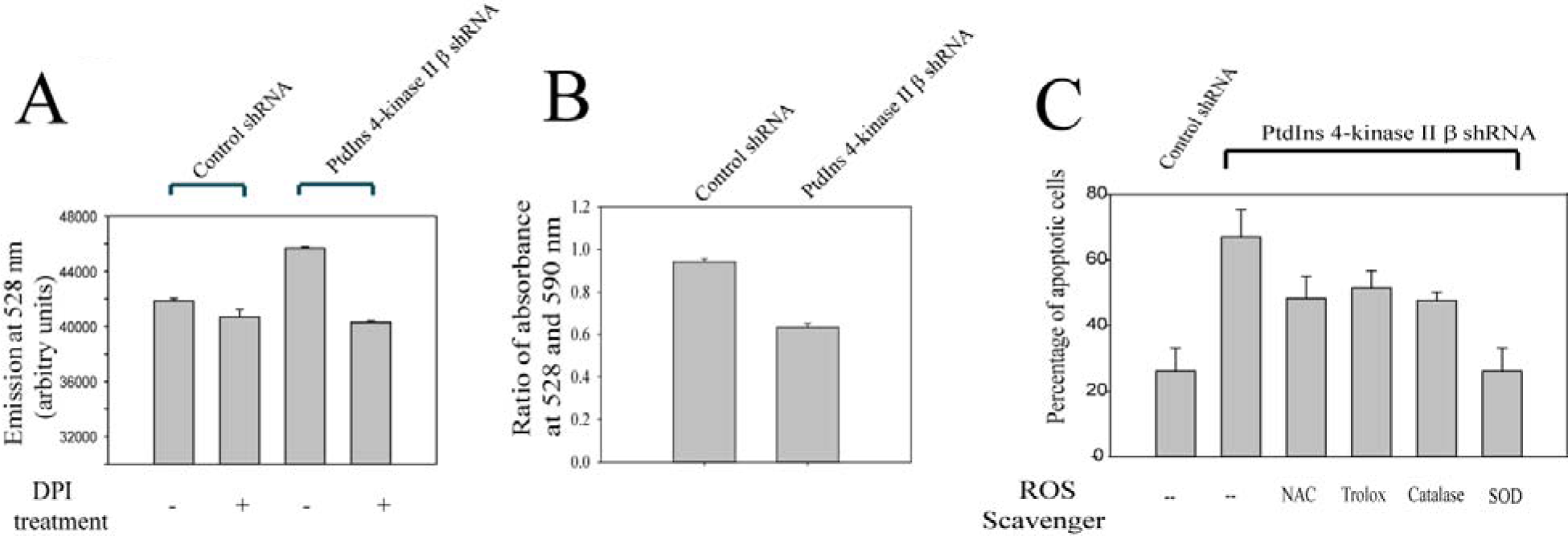
**A. Diphenyleneidonium (DPI) abrogates increased ROS levels in PtdIns 4-kinase II β depleted cells.** MCF-7 cells were incubated with CM-H_2_DCFDA and DPI (50µM) for 1 h. These cells were transfected with either control or PtdIns 4-kinase II β shRNA plasmids and incubated for 24 h. At the end of incubation, fluorescence was measured (excitation: 485 nm, emission: 528 nm and gain: auto scale) using microplate reader. **B. Mitochondrial depolarisation**. After 24 h of PtdIns 4-kinase II βshRNA transfection, loss of mitochondrial membrane potential was estimated using JC-1 dye. **C. Effect of ROS scavenger on PtdIns 4-kinase II** β **induced cell death**. MCF-7 cells were pre-incubated with NAC (10 mM), Trolox (1 mM), SOD (135 U/ml) and catalase (200 U/ml) for 1 h. These cells were transfected with either control or PtdIns 4-kinase II β shRNA plasmids and incubated for further 36 h. These cells were assayed for apoptosis using propidium iodide. (The data indicates mean ± SD (P<0.001). The experiment were repeated thrice)

Pre-incubation with thiol dependent (N-acetyl cysteine and trolox) and independent (PEG-superoxide dismutase and PEG-catalase) ROS scavengers, protected cells from PtdIns 4-kinase II β shRNA induced apoptosis (Fig. 2C). Superoxide dismutase was most effective treatment with almost 100% reversal. Other scavengers showed reversal in cell death with varying efficiency: SOD> NAC> catalase>trolox.

### Depletion of PtdIns 4-kinase II β inhibits NF-κB activity and suppresses pro-survival gene expression

In earlier studies from our lab, PtdIns 4-kinase II β knockdown was shown to stabilize Par-4 protein levels and facilitates nuclear translocation. In nucleus, Par-4 levels were shown to inhibit NF-κB binding to its promoter sequence and suppress Bcl-2 and inflammatory responsive genes expressions ^23^. Nuclear extracts from PtdIns 4-kinase II β shRNA transfected cells showed a reduction in NF-κB binding to its target sequences. Super shift assay with antibodies against NF-κB subunit, p65 showed low levels of NF-κB in nuclear extracts of PtdIns 4-kinase II β depleted cells (Fig. 3A).Inhibition of NF-κB may alter redox homeostasis and suppresses prosurvival genes, Bcl-2 and Akt-3 ^9,24^. The effect of PtdIns 4-kinase II β shRNA transfection on NF-κB regulated pro-survival genes, Bcl-2 and Akt-3 expression was studied. PtdIns 4-kinase II β knockdown showed reduction in both Bcl-2 and Akt-3 mRNA levels in comparison to control cells (Fig. 3 B).

**Figure 3.**
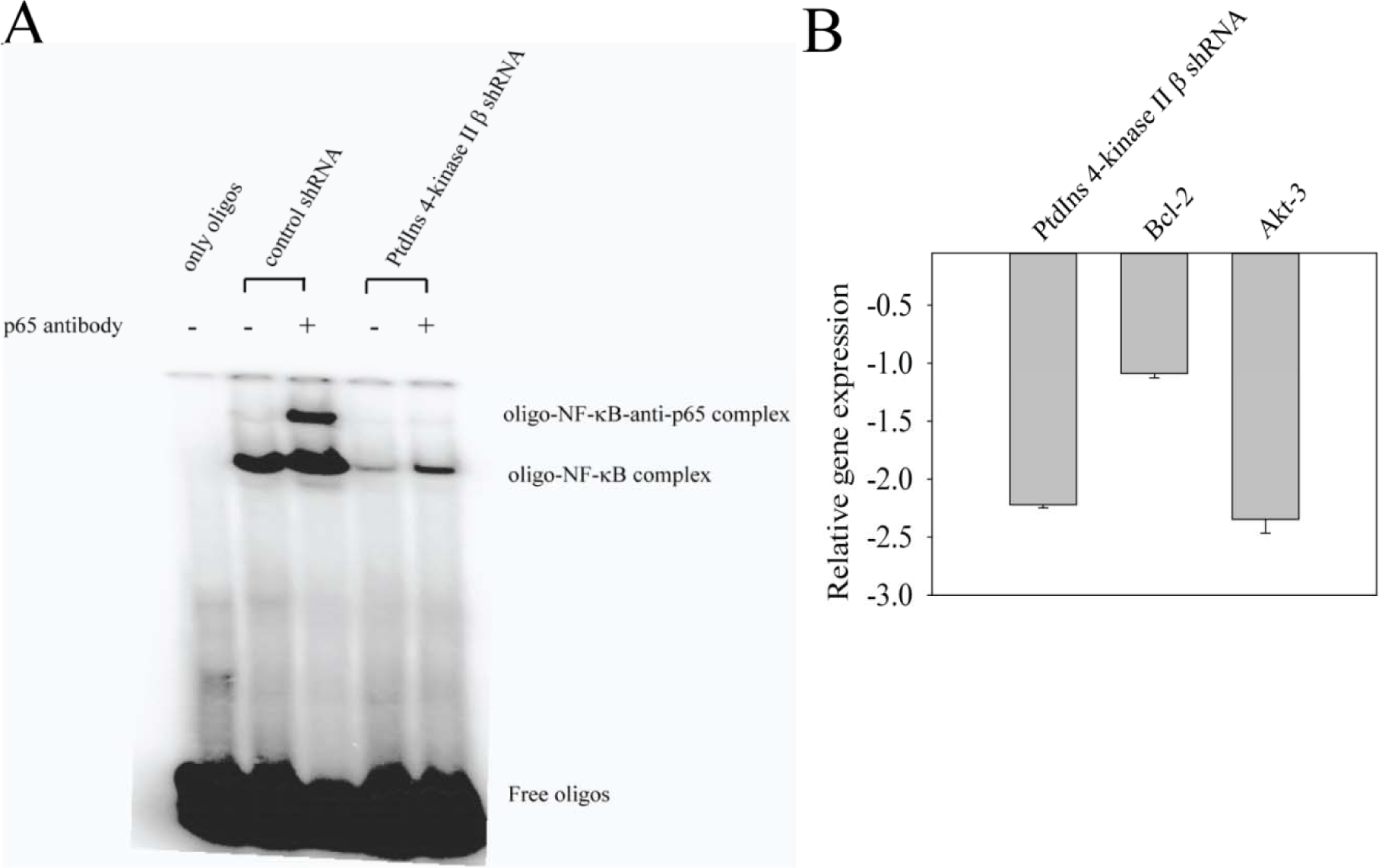
PtdIns 4-kinase II β shRNA affects NF-κB activity. **A**.MCF-7 cells were transfected with either control or PtdIns 4-kinase II β shRNA plasmids and incubated for 24 h. Nuclear extracts were incubated with end labelled NF-κB specific oligonucleotides and antibodies against p65. The complex was electrophoresed as described in Materials and Methods. **B**. MCF-7 cells were transfected with either control or PtdIns 4-kinase II β shRNA plasmids and incubated for 24 h. PtdIns 4-kinase II β, Bcl-2 and Akt-3 mRNA levels were quantified using gene specific primers. GAPDH was used for normalization. The data indicates mean ± SD of three independent experiments (P<0.001).

### PtdIns 4-kinase II β depletion correlates with increased levels of ER stress

Oxidative stress results in misfolding of proteins which leads to ER stress. Further, accumulation of misfolded or unfolded proteins in ER lumen leads to activation of unfolded protein responses (UPR), resulting in activation of pro survival as well as apoptotic pathways. PtdIns 4-kinase II β depletion resulted in increased levels of ER stress markers, PERK, IRE-1α, Bip, Ero1-Lα, calnexin and PDI proteins (Fig. 4A). Protein levels of Bip and PDI showed a profound increase in PtdIns 4-kinase II β shRNA transfectants.

**Fig. 4.**
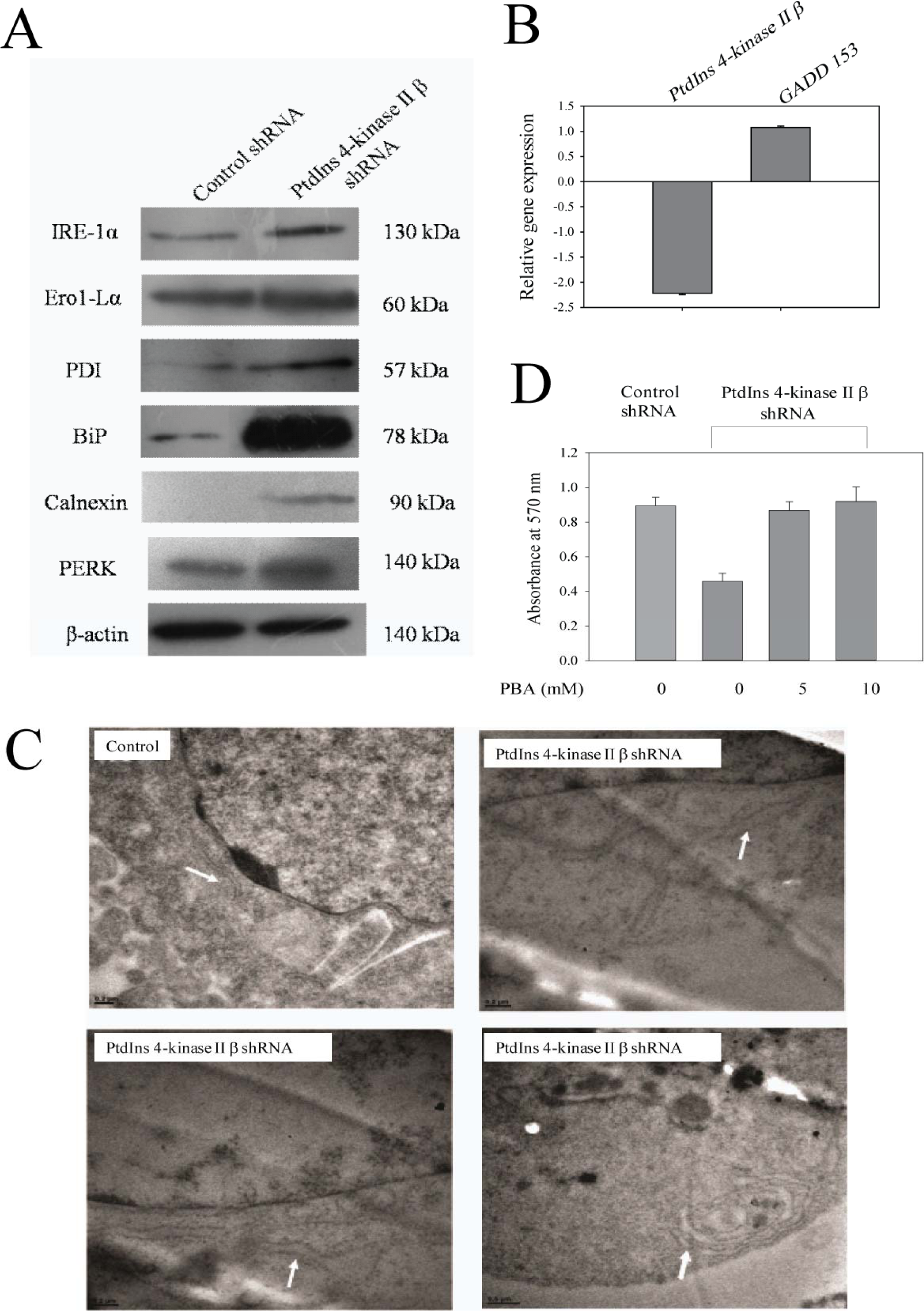
PtdIns 4-kinase II β depletion induces ER stress in MCF-7 cells. **A**.MCF-7 cells were transfected with either control or PtdIns 4-kinase II β shRNA plasmids and incubated for 36 h. Cell lysates were analysed for ER stress marker proteins using antibodies. β-actin was used as loading control.**B**.Relative expression of GADD153. **C**. Transmission electron micrographs of control and PtdIns 4-kinase II β shRNA transfected MCF-7 cells. Arrow indicates ER.**D**.MCF-7 cells were incubated with PBA for 1 h and transfected with either control or PtdIns 4-kinase II β shRNA plasmids. After 36 h of incubation, cell viability was determined by MTT assay. The data indicates mean ± SD (P<0.001). The experiment was repeated thrice.

Quantitative PCR analysis showed an increase in GADD153 gene expression in PtdIns 4-kinase II β shRNA transfected cells within 24 h (Fig. 4B). GADD153 up regulation was correlated with down regulation of Bcl-2 and Akt-3.Transmission electron micrographs of PtdIns 4-kinase II β shRNA transfectants showed an increase in ER lumen diameter in comparison to control cells. These micrographs indicate unresolved ER stress that leads to accumulation of unfolded proteins in ER lumen resulting in dilation of ER. ER lumen deformities provide further evidence for ER stress in PtdIns 4-kinase II β depleted MCF-7 cells (Fig. 4C).

4-phenyl butyric acid (PBA), a chemical chaperone, is shown to protect cells from ER stress. Pre incubation with PBA (5 mM) rescued MCF-7 cells from PtdIns 4-kinase II β shRNA induced cell death with almost 100% efficiency (Fig. 4D).These studies indicated that PtdIns 4-kinase II β depleted cells undergo ER stress.

### PtdIns 4-kinase II β depletion increases radio sensitivity of MCF-7 cells

Redox status of the cell plays an important role in radiation sensitivity in cancer cells. Irradiated MCF-7 cells showed dose dependent increase in levels of PtdIns 4-kinase II β mRNA (Fig. 5A). U87-MG cells (gliobalstoma cell line) also showed increased PtdIns 4-kinase II β mRNA expression upon radiation and radioresistant U87-MG cells showed higher expression levels of PtdIns 4-kinase II βmRNA as compared to radiosenstitive U87 MG cells (results not shown).

**Figure 5.**
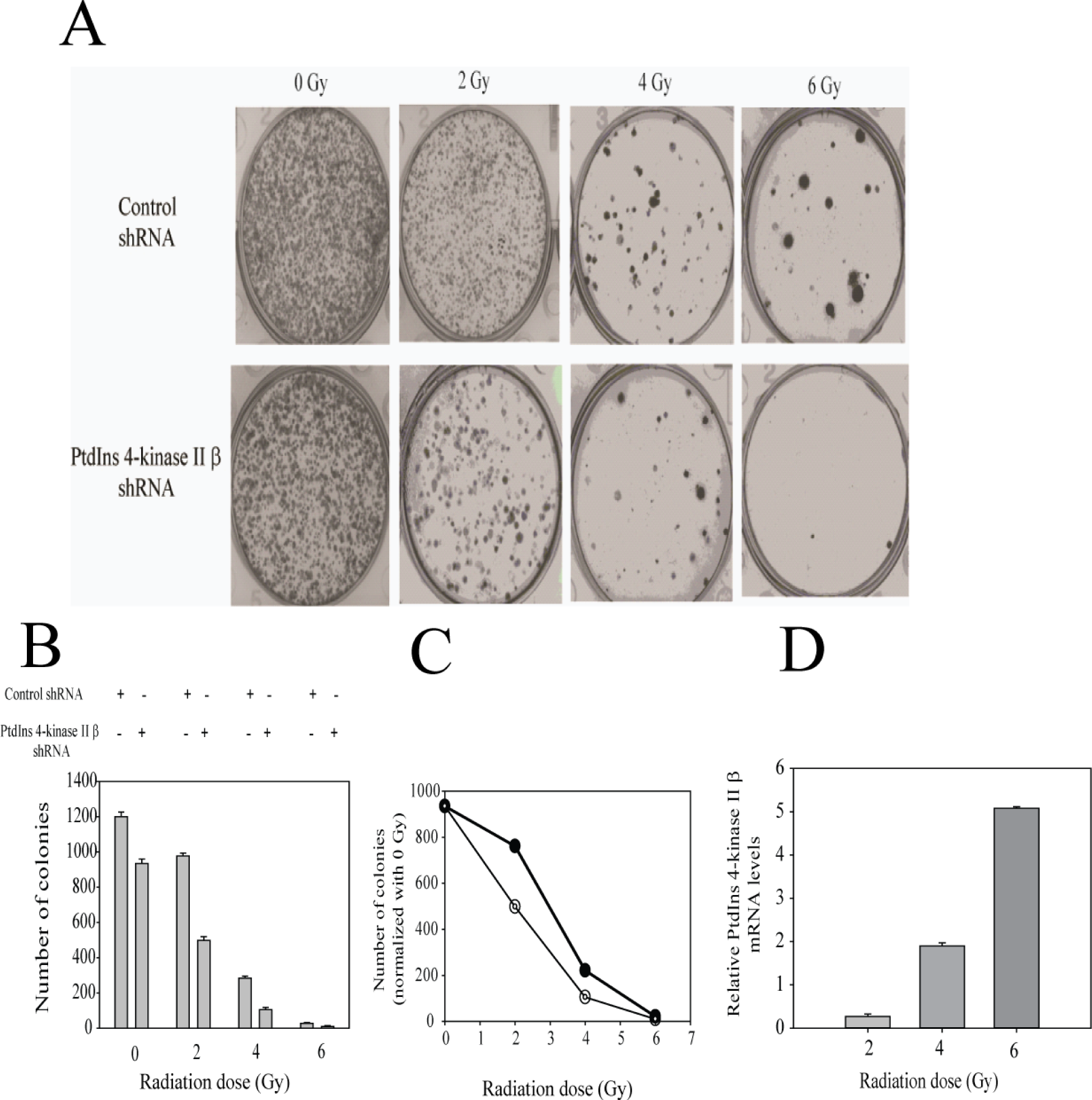
PtdIns 4-kinase II β depletion increases radio sensitivity of MCF-7 cells. MCF-7 cells were transfected with either control or PtdIns 4-kinase II β shRNA plasmids and incubated for 12 h. After incubation, 2000 cells were plated for clonogenic assay and irradiated with indicated radiation dose. **A)** Representatives images of colonies and numbers on the top of images indicate radiation dosage. **B)** Quantification of the clonogenic data. The bar graph represents mean ± SD (P<0.001). **C)** Survival curve with number of colonies normalized with 0 Gy (○-○ control shRNA, ○-○ PtdIns 4-kinase II β shRNA). The experiment was repeated thrice. D) MCF-7 cells were irradiated with indicated dose. After 24 h, PtdIns 4-kinase II β mRNA levels were quantified using quantitative PCR.

Further, increased doses of γ-radiation showed reduction in the colony forming ability of MCF-7 cells. Approximately 20% reduction in colony forming ability was observed with 2 Gy radiations and 80% reduction was observed with 4 Gy radiations in control cells. However, PtdIns 4-kinase II β depleted cells showed 48% less colonies after2 Gy radiations comparison to control cells. At higher doses (4 Gy and 6 Gy), the total number of colonies were very few making it difficult for any statistical inference (Fig. 5B). These results suggest that PtdIns 4-kinase II β depletion increases radio sensitivity of malignant cells. These results can have potential implication in overcoming radio-resistance of cancer and improving the outcome of radiotherapy.

## DISCUSSION

Knockdown of PtdIns 4-kinase II β has earlier been shown to stabilize Par-4 levels and facilitates Par-4 nuclear localization ^7^. Par-4 affects stress signaling pathways mediated by NF-κB, transcriptional factor and inhibits NF-κB activity through different mechanisms in cytoplasm as well as in nucleus. In cytoplasm, Par-4 inhibits PKC ζ mediated phosphorylation of IκB and NF-κB nuclear translocation ^25^. In the nucleus, over expression of Par-4 inhibits NF-κB binding to its promoter sequences ^23^. Inhibition of NF-κB also alters redox homeostasis and suppresses the expression of prosurvival genes, Bcl-2 and Akt-3 ^9,24^. In the present study we observed that PtdIns 4-kinase II β depletion resulted in reduction in nuclear levels of NF-κB probably mediated through Par-4. In addition, PtdIns 4-kinase II β depletion correlated with reduced expression of Bcl-2 and Akt-3, the downstream targets of NF-κB. Bcl-2 is localized to outer mitochondrial membrane. It inhibits the actions of pro-apoptotic proteins, such as Bak and Bax. These proapoptotic proteins are involved in mitochondrial membrane pore formation. Decreased Bcl-2 expression is correlated with increased production of ROS.

NF-κB also modulates ROS levels in cells by regulating expression of superoxide dismutase, ferritin heavy chain and glutathione S-transferase genes ^13^. Under normal conditions, the molar ratio of GSH and GSSG exceeds 100:1. However, when cells were exposed to oxidative stress this ratio decreases to 10:1 and even 1:1 ^26^. Reduction in GSH: GSSG ratio to 6:1 and increased ROS levels in PtdIns 4-kinase II β depleted cells suggest induction of oxidative stress in these cells. This hypothesis was further supported by the observation that ROS scavengers were able to rescue MCF-7 cells from PtdIns 4-kinase II β shRNA induced apoptosis.

ROS is produced in cells either as a byproduct of biological processes or by NADPH oxidases as part of their biological function.NADPH oxidase inhibitor abrogated increased ROS levels in PtdIns 4-kinase II β shRNA transfectants indicating a role of NADPH oxidases. In a contrasting report, antibodies against type II PtdIns 4-kinase have been shown to significantly reduce oxidative burst in N-formyl-methionyl-leucyl-phenylalanine stimulated neutrophils ^27^. This variation may be due to various isoforms of NADPH oxidase that may be regulated differently. In phagocytic cells, Nox2 generates oxidative bursts, important in host defense responses. Nox5 has been demonstrated in many non-immune cells and tissues including cancer cells. NF-κB has also been shown to regulate Nox5 in human aortic smooth muscle cells ^28^. More studies are needed to identify the NOX regulated by PtdIns 4-kinase II β.

Oxidative stress induction through NADPH oxidase activation plays an important part in ER stress induced apoptosis^29^. Earlier, the immuno-proteomic data of PtdIns 4-kinase II β showed presence of ER associated proteins as interacting partners, suggesting that PtdIns 4-kinase II β may be involved in ER associated functions^7^. Increased levels of PERK, IRE-1α, Ero1-Lα, calnexin, Bip, and PDI isomerase in PtdIns 4-kinase II β depleted cells suggest induction of UPR in these cells. Apart from mediating isomerisation of disulfide bonds during protein folding, PDIA5 has also been suggested to cleave disulfide bonds in ATF-6. Active ATF-6 translocates to nucleus and up regulates transcription of ER chaperones such as Bip^30^. PDIA5 depleted cells do not undergo oligomer dissociation, ER-to-Golgi migration and induction of ATF6 target genes ^31^. Shot gun proteomic approach has shown the presence of PDIA5 in PtdIns 4-kinase II β immunoprecipitates ^7^. The mechanisms involved in PtdIns 4-kinase II β and PDIA5 interactions are not known at present and need further studies. During ER stress, increased load of mis-folded proteins clog ER which results in deformities in ER structure. Presence of dilated ER lumen in PtdIns 4-kinase II β depleted cells supported this hypothesis. The ability of PBA, chemical chaperone, to rescue cells from PtdIns 4-kinase II β shRNA induced ER stress further supported accumulation of misfolded proteins in ER lumen.The mechanism by which PtdIns 4-kinase II βinduced ER stress needs investigation.

Radiotherapy is one of the important segments in cancer treatment. One of the limiting factors of radiotherapy is cytotoxicity to the normal cells. Combinatorial therapy with other drugs might help in decreasing the dose required as well as reducing cytotoxicity to normal cells. We observed PtdIns 4-kinase II β depletion induced radio sensitivity in MCF-7 cells. The increase inPtdIns 4-kinase II β mRNA expression in irradiated MCF-7 cells indicated the potential role of PtdIns 4-kinase II β over expression in rendering cancer cells radio resistant.

These studies indicate that PtdIns 4-kinase II β depletion results in inhibition of NF-κB nuclear translocation and induction of oxidative stress as well as ER stress culminating in cell death. These results have potential implication in radiation therapy for management of radio-resistant cancers.

## Notes

### Competing Interest Statement

The authors have declared no competing interest.

